# Modeling and Detecting Downward Causation in Disease Transmission

**DOI:** 10.64898/2026.10.20.760281

**Authors:** Meilin Yen, Pranav Muralikrishnan, John Harte

## Abstract

We compare two epidemic models: the deterministic SIR (Susceptible-Infected-Recovered) model and the maximum-entropy-based DyMES (Dynamic Maxent across Entwined Scales) framework. In contrast to the widely used SIR model, the recently proposed DyMES model incorporates both top-down and bottom-up mechanisms and thus can capture local responses to global behavior. We simulate both models in three disease scenarios using parameter values drawn from the literature and analyze trends in the epidemic trajectories. Increasing the strength of downward causation lowers the peak but extends the time taken to return to the initial state, changing the trajectory from an SIR-like rise and fall to an epidemic with a long tail. By combining the Maxent inference procedure and top-down causation, the DyMES model predicts the probability distribution of disease incidence across population sub-aggregates.

## 1. Introduction

Epidemiological models are key tools for understanding and combating epidemics. These models help us predict the infection trajectories of epidemics and the consequences of remedial strategies. The compartmental SIR (Susceptible-Infected-Recovered) model is one such epidemiological model. Under this model, individuals in the system move between susceptible (*S*), infected (*I*), and recovered (*R*) states, or “compartments.” Time-differential equations, one per compartment, govern these movements [1]. The simplicity of this version of the SIR framework makes it a good starting point for infectious disease modeling. However, this model tends to oversimplify disease transmission, ignoring, for example, the effect of disseminated information about system-wide disease incidence on individual behavior within groups. This oversimplification can lead to inaccurate predictions.

Using a more mechanistic framework can help us avoid some of these limitations. DyMES (Dynamic Maxent across Entwined Scales) is a recently developed theory describing the dynamics of complex systems in which causal influence flows bidirectionally between higher and lower levels of organization [2, 3]. Here, Maxent refers to the principle of maximizing Shannon information entropy under constraints imposed by macrovariables. The theory predicts both the time-evolving probability distribution over the microvariables and the trajectories of the macrovariables. In our application of DyMES, top-down causation arises because the rate constant governing disease transmission between individuals is influenced by the incidence of the disease in the larger population if individuals heed public health warnings when that disease incidence is high [3]. In applying the DyMES model to a real-world system, we want to explore the model’s strengths and weaknesses. Additionally, seeing where and how the DyMES framework differs from other approaches to modeling infectious diseases helps clarify DyMES’s potential role within current epidemiological research.

Here, we compare the output of a deterministic SIR model and an epidemic-adapted version of the Maxent-based DyMES model in simulating three hypothetical epidemic scenarios, each corresponding to a real-world disease. By comparing the results of these scenarios, we can begin to assess the importance of, and capacity to detect, downward causation as a potentially important factor influencing the spread of disease.

## 2. Models

### 2.1. Shared assumptions

In both our DyMES and SIR models, we consider multiple disjoint populations of individuals in some region. The disease spreads between individuals in the same group; an infected individual in one group cannot transmit the disease to a susceptible individual in another. Infection spreads via contact with infected individuals, rather than from environmental contact or sexual transmission. Individuals cannot travel between groups. We also assume that the strain of disease present in the system does not mutate throughout the course of our simulation. We ignore birth, natural death, and reinfection (i.e., if individuals recover instead of dying, they cannot contract the disease again).

The DyMES and SIR models share the following parameters: *G* (the number of groups), *c*_0_ (the transmission rate), and *d*_0_ (the death/recovery rate per day). For both models, *G* = 100. There are 100 individuals in each group and 100 total infected individuals at time *t* = 0. These parameters are presented in Table 1.

**Table 1.**
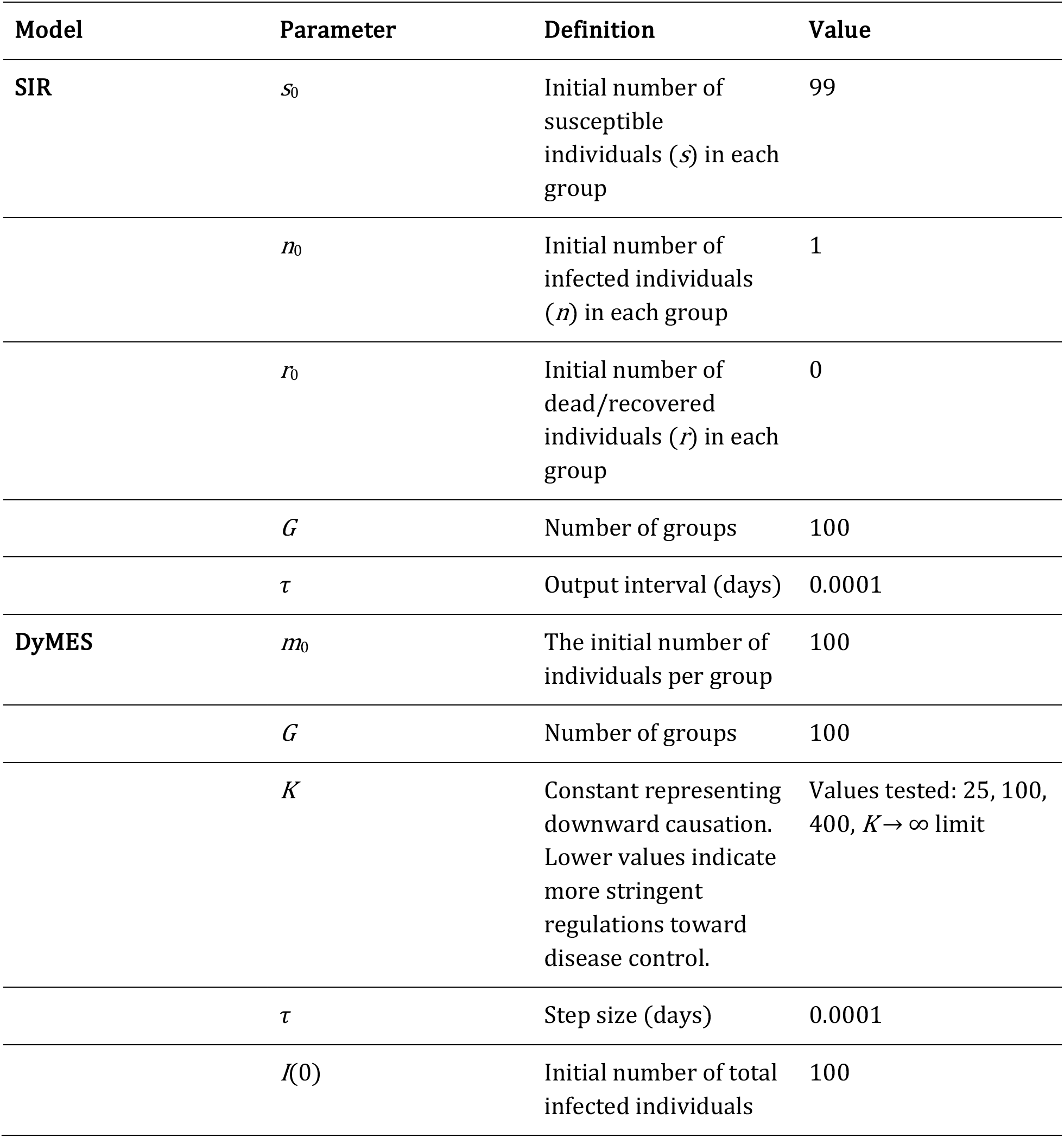
The constants in the SIR and DyMES models, their definitions, and the values used in the simulations.

We examine three different epidemic scenarios: (1) high *c*_0_ and high *d*_0_, (2) medium *c*_0_ and medium *d*_0_, and (3) low *c*_0_ and low *d*_0_. The scenarios correspond loosely to the following real-world epidemic systems: (1) highly pathogenic avian influenza (*Alphainfluenzavirus influenzae*) (HPAI) H5N1 in chickens (*Gallus gallus domesticus*), (2) Newcastle disease virus (*Orthoavulavirus javaense*) (NDV) in commercial broiler chickens, and (3) foot-and-mouth disease (*Aphthovirus vesiculae*) (FMD) in dairy cows (*Bos taurus*). These diseases were chosen because they fit the shared assumptions. Different serotypes of FMD cause similar infections, so the specific serotype in the system does not matter [4]. The parameters are presented in Table 2.

**Table 2.** The different values of *c*_0_ and *d*_0_ tested in the three cases.

| Disease | $c_0$ (new infections per susceptible-infected pair per day) | $d_0$ (number of deaths/recoveries per infected individual per day) | Data Source |
| --- | --- | --- | --- |
| HPAI (high $c_0$/high $d_0$) | 0.0123 | 0.5 | [5, 6] |
| NDV (medium $c_0$/medium $d_0$) | 0.0067 | 0.2 | [7] |
| FMD (low $c_0$/low $d_0$) | 0.0033 | 0.167 (1/6) | [8] |

### 2.2. SIR model

Each group in our SIR system has three compartments: individuals susceptible to the disease (*s*), infected individuals (*n*), and recovered or deceased individuals (*r*). After calculating the trajectories for one group, we multiply the numbers of susceptible and infected individuals by *G* to predict infection rates across all *G* groups. We assume that all groups behave identically and that all groups start with one infected individual per group.

While the classical SIR model assumes identical behavior across all groups, we introduce this group notation because we will be comparing its output against the DyMES model, which predicts a probability distribution for the number of infections at any moment in time across groups.

The differential equations for one group, for constant *T* = *s* + *n* + *r* and definitions as described in Tables 1 and 2, are defined as follows:

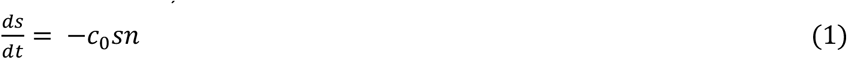

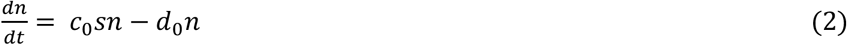

The differential equation for *r* follows naturally from Eqs. 1 and 2.

### 2.3. DyMES model

Many complex systems exhibit both top-down and bottom-up causation, thereby entwining the macro-level and micro-level variables. DyMES is a dynamic theory of such systems; it predicts both the time-evolving probability distributions over microvariables and the time evolution of the macrovariables. DyMES utilizes the principle of maximum entropy (Maxent) to calculate, at every instant, microvariable distributions subject to constraints imposed by the macrovariables and their first time derivatives. The transition functions, a key feature of DyMES, encode the dependence of the time derivatives of the mean values of microvariables on both micro- and macrovariables, where the latter dependence captures top-down causal influence [3].

Our DyMES model describes a population of individuals separated across geographically disjoint groups. An infected individual in one group can spread the disease to any other uninfected individual in that group. However, that individual cannot transmit the disease to uninfected individuals in other groups. Our system contains implicit spatial structure, but we do not explicitly encode any spatial components in our model. The groups can behave heterogeneously in the DyMES model, which contains a probability distribution over the microvariable. We take expectations over this distribution to encapsulate average behavior.

In our model, downward causation is represented by preventative measures that affect the transmission rate. Some potential examples include increases in the administration of therapeutic measures that reduce transmission, hand washing, disinfecting of group spaces, and minimizing the transport of items inside and outside each group. Levels of downward causation change in response to information on the whole system, such as news updates from central authorities. Different models have implemented vaccination as a mechanism that decreases the pool of susceptibles and/or reduces the transmission rate [5]. We assume that, in our system, vaccination as a mechanism of downward causation acts on the transmission rate rather than the susceptible pool.

The microvariable in this system is *n*(*t*), the number of infected individuals in a group at time *t*. The macrovariable is *I*(*t*) = *G*⟨*n*⟩, the total number of infected individuals in the system at time *t*. For this microvariable-macrovariable pair, the time-dependent probability distribution *P*(*n*) corresponds to the probability that there are *n* infected individuals in a group. At time *t*, let the expectation of the distribution be ⟨*n*⟩_*t*_. Here, *m*_*t*_ is the total number of alive, non-recovered individuals in each group at time *t*. At the start of the simulation, *m*_0_ = 100 and ⟨*n*⟩_0_ = *I*(0) / *G* = 1.

The two constraints in this application of DyMES are:

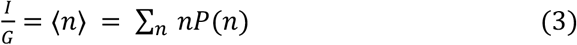

and

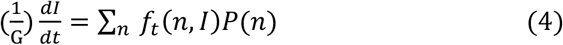

*f*_*t*_, the transition function, is written as:

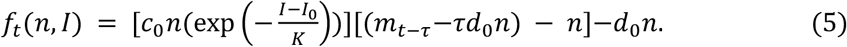

Here, *m*_*t*_ is updated at every timestep as *m*_*t*_ = *m*_*t*−τ_−*τd*_0_*n*. When we subtract *n* from *m*_*t*_, the resulting term is analogous to the variable *s* in Eq. 2. Multiplying *m*_*t*_ − *n* by 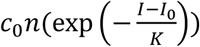 yields the increase in infected individuals due to mixing between infected and susceptible (i.e., uninfected and alive) individuals. The parameter *τ* is the step size in days. The −*τd*_0_*n* term is the number of removed (i.e., dead or recovered) individuals from the total pool of individuals per timestep, and the −*n* term further removes infected individuals from the total pool of alive (and non-recovered) individuals. When *t* = 0, we use *m*_0_ instead of *m*_*t*−τ_.

These two terms allow the number of susceptible individuals to decrease as time passes in the simulation. The −*d*_0_*n* term, which is analogous to the second term on the right-hand side of Eq. 2, represents the decrease in the number of infected individuals due to death or recovery from disease.

Downward causation is represented phenomenologically by the 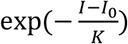 term in Eq. 5 and only comes into effect once disease incidence increases past the starting *I*_0_. When *I* < *I*_0_, this term no longer makes sense, as 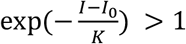. Therefore, we only look at the DyMES model’s behavior from the start of the epidemic to when the number of infected individuals returns to *I*_0_. We tested *K* = 25, 100, and 400, as well as the *K* → ∞ limit, which corresponds to no downward causation.

After maximizing the information entropy −*Σ*_*n*_*P*(*n*)log(*P*(*n*)) of *P*(*n*) under these constraints, the probability distribution 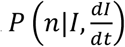 is given by:

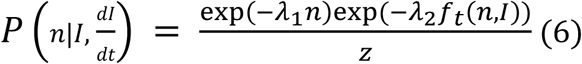

where *z* is a normalization constant and the *λ*_*i*_ are Lagrange multipliers. DyMES is solved numerically with an iterative procedure. At timestep *i* the probability distributions are constrained by the macrovariables and their time derivatives. The macrovariables are advanced to step *i* + 1 using their known time derivatives at step *i*. To update the time derivatives of the macrovariables, the updated macrovariables are substituted into the transition functions, which are then averaged over the distribution evaluated with the step *i* Lagrange multipliers. Thus, over multiple iterations of DyMES, these steps form a feedback loop linking time-dependent microvariable probability distributions with macrovariable dynamics. For more theoretical detail and a worked-out example, see [3].

The DyMES parameter values are presented in Tables 1 and 2.

## 3. Methods

To find the parameters of our model, we searched the literature for past epidemic modeling studies that were similar to our DyMES model. This model requires that groups be homogeneous and mutually exclusive and that disease cannot spread between two individuals in two different groups. Given these requirements, we focused our literature search on livestock epidemics. Livestock animals are relatively homogeneous (compared to humans). They cannot travel between farms, meaning disease cannot be spread between individuals in different groups. Additionally, one individual cannot be part of two farms. We also focused our search on livestock diseases that spread via non-sexual contact between infected and uninfected individuals rather than environmental, vertical, or sexual transmission.

We implemented the SIR and DyMES models using Python. For the SIR model, we used scipy.integrate.solve_ivp with the adaptive Runge–Kutta order 5(4) method [9, 10] and stored solutions every *τ* days.

For the DyMES model, we used a modified version of the code published in [3]. The modified version calculates expectations over a probability distribution from 0 to *m*_*t*−τ_, rather than from 0 to the macrovariable *I*. This change is necessary because the number of infected individuals in any one group cannot exceed *m*_*t*−τ_, which decreases over time. The code also stops the model from simulating once the macrovariable *I* decreases to below its starting value (*I*_0_). We used scipy.optimize.fsolve to find the Lagrange multipliers that satisfy the constraints. For each simulation, we started with a value of *τ* = 0.0001.

We ran each DyMES model until the macrovariable returned to its initial state. Then, we ran the SIR model for at least as long as required for the SIR trajectory to return to 100. For any runs in which the DyMES constraints were no longer satisfied past a certain point, we only considered results up to just before that point.

## 4. Results

All the disease incidence curves displayed the same overall trend: disease incidence rose, peaked, and then fell at a slower rate than the initial rise. Across all tested cases, decreasing *K* in the DyMES model noticeably reduced disease spread (Figure 1, Table A.1). Lower values of *K* mean increased downward causation. When *K* decreases, 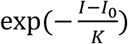 decreases for any given value of *I* > *I*_0_, lowering the effective transmission rate as well. As *K* decreased and downward causation increased, the curve flattened, the peak was smaller and earlier in the epidemic, and the disease incidence fell over a longer timescale. The pattern of a fast initial peak and a longer drop back to 100 was most visible for small *K*, and increasing *K* did not seem to increase the peak size or decrease the time taken to return to *I* = 100 by a linear factor.

**Figure 1:**
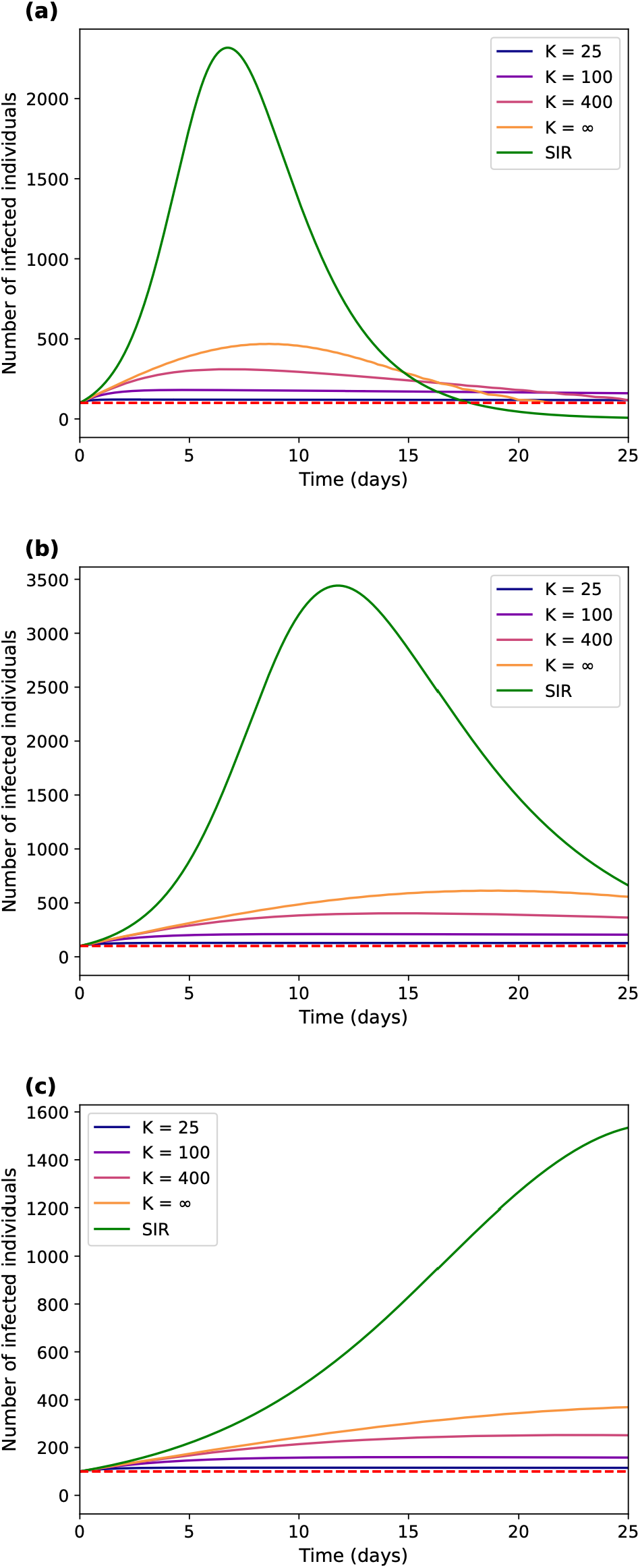
Infection trajectories for DyMES and SIR. These graphs display the numbers of infected individuals *I*(*t*) for the first 25 days for the finite-*K* DyMES models, the DyMES model at the *K* → ∞ limit, and the SIR model for (a) HPAI (high *c*_0_/high *d*_0_), (b) NDV (medium *c*_0_/medium *d*_0_), and (c) FMD (low *c*_0_/low *d*_0_). The red dashed line represents *I* = 100.

Across the DyMES graphs, lower values of *d*_0_ were generally associated with increased time taken to return to *I* = 100. When we compared the results of the high and low *c*_0_/*d*_0_ cases, we saw that disease incidence took longer to return to 100 in the *K* → ∞ limit for the low *d*_0_ case compared to the high *d*_0_ case. Because the *K* → ∞ limit model constraints could no longer be satisfied in the medium *c*_0_/medium *d*_0_ case after *t* = 46.91, we only have a lower bound for the return time. Our lower bound partly supports this finding, as it is greater than the return time for the high *c*_0_/high *d*_0_ case (Table 3), but further testing is necessary to confirm that it is also less than the return time for the low *c*_0_/low *d*_0_ case. Higher *c*_0_ was also associated with decreased time to reach the peak.

**Table 3.** Times taken to reach peak disease incidence and to return to 100 for the DyMES *K* → ∞ limit and SIR models across all three cases. Lower bounds exist where exact return times cannot be calculated due to constraints not being satisfied.

| Disease | DyMES $K \rightarrow \infty$ limit time to reach peak (days) | DyMES $K \rightarrow \infty$ limit time to return to 100 (days) | SIR time to reach peak (days) | SIR time to return to 100 (days) |
| --- | --- | --- | --- | --- |
| HPAI (high $c_0$/high $d_0$) | 8.63 | 21.46 | 6.75 | 17.78 |
| NDV (medium $c_0$/medium $d_0$) | 19.05 | > 46.91 | 11.78 | 36.09 |
| FMD (low $c_0$/low $d_0$) | 27.82 | 63.85 | 26.62 | 64.40 |

However, a higher *c*_0_ did not guarantee a higher peak, as the peak for the medium *c*_0_/medium *d*_0_ case was higher than that of the high *c*_0_/high *d*_0_ case.

Higher values of *K* resulted in larger deviations in the DyMES probability distributions from the associated geometric distribution with mean ⟨*n*⟩, which is the distribution that maximizes entropy for some mean ⟨*n*⟩ in the Maxent framework [11, 12]. The DyMES distribution’s deviation from the geometric distribution introduces curvature into the log probability distribution via the *n*^2^ terms in Eq. 5. As *I*(*t*) increased, probability mass shifted toward higher possible values and the curve flattened. Changing the values of *c*_0_ and *d*_0_ also appeared to affect the curvature across the same values of *K*. All these trends were especially visible for large *K* (Figure 2).

**Figure 2:**
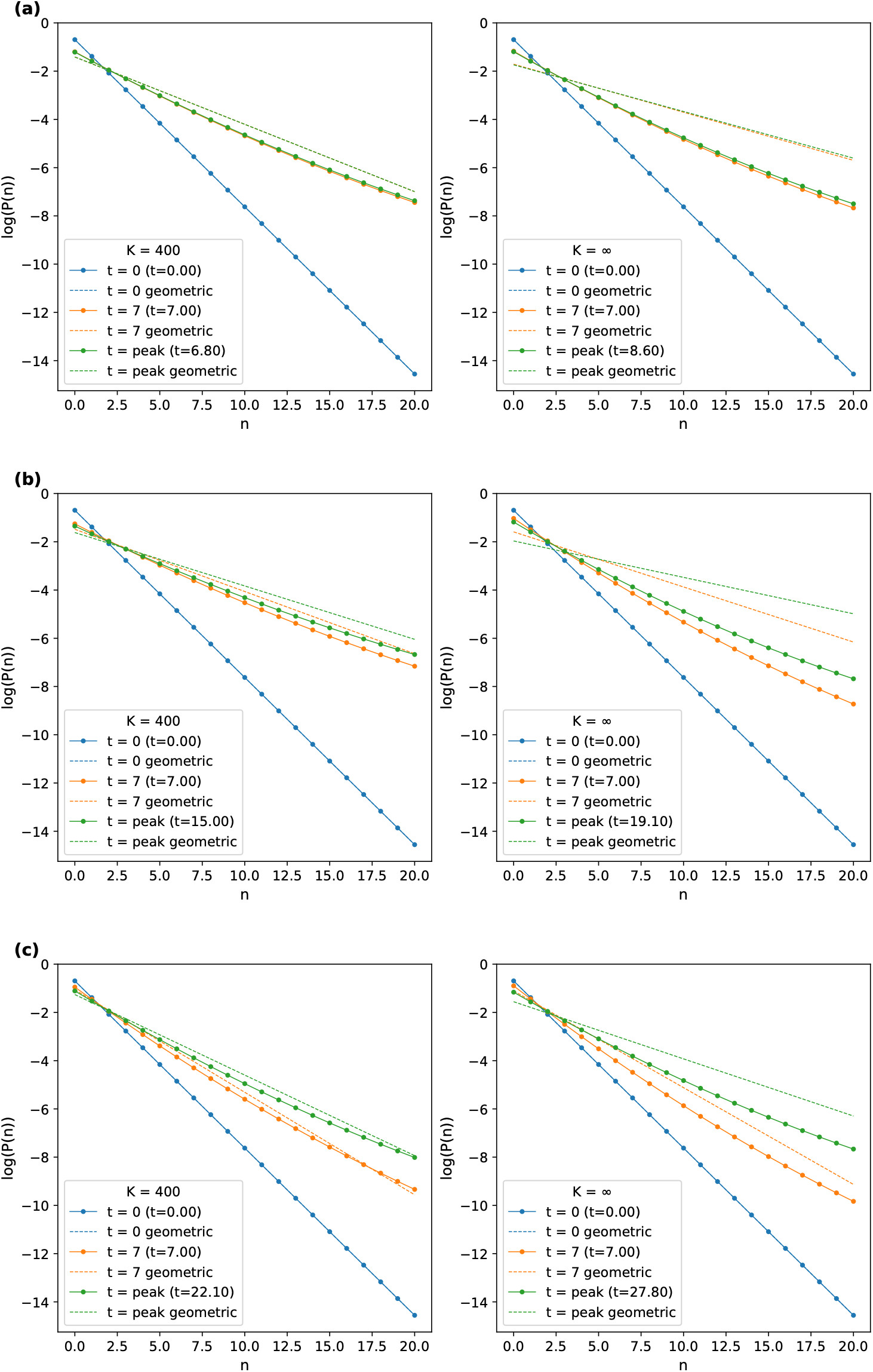
Log probability distributions for DyMES and geometric distributions with the same. ⟨***n***⟩. Natural log probability distributions for the DyMES models with high *K* (*K* = 400, *K* → ∞ limit) and the geometric distributions with the same ⟨*n*⟩ at times *t* = 0, *t* = 7, and *t* = peak (rounded to the nearest tenth), for (a) high *c*_0_/*d*_0_, (b) medium *c*_0_/*d*_0_, and (c) low *c*_0_/*d*_0_. *K* = 25 and *K* = 100 were omitted from the figure because the high levels of downward causation made the differences in the distributions negligible.

Unlike the DyMES models, the SIR model cannot capture the long-tail phenomenon, as there is no way to incorporate downward causation imposed by finite *K* into the model. Compared to the DyMES model at the *K* → ∞ limit, the SIR model’s peaks were earlier and considerably higher (Table 3, Figure 1). In the cases with high and medium *c*_0_, the *K* → ∞ limit reached 100 after the SIR model. This relationship was flipped in the low *c*_0_ case (Table 3). When *c*_0_ and *d*_0_ were low, the *K* → ∞ limit reached 100 just before the SIR model. Across both the DyMES and SIR models, disease incidence took the longest to peak in the FMD case (with low *c*_0_ and low *d*_0_).

For all runs in the high and low *c*_0_ and *d*_0_ cases, the percent error of *G*⟨*n*⟩ relative to *I* never exceeded 1%, except for a single timestep at *t* = 19.98 in the high *c*_0_/high *d*_0_ case for the *K* → ∞ limit. At this time, the solver briefly failed to converge (percent error = 83%) but quickly recovered. The errors were low throughout for the DyMES *K* = 25 and *K* = 100 models with medium *c*_0_/medium *d*_0_. For *K* = 400 and the *K* → ∞ limit with medium *c*_0_/medium *d*_0_, the errors were similarly small until *t* = 61.53 and *t* = 46.91, respectively. After these times, Eq. C.2 was no longer satisfied, and the Lagrange multipliers and percent errors grew rapidly. When we decreased *τ* by a factor of 10 (*τ* = 0.00001), the constraints failed at the same times, indicating that this failure was due to model behavior rather than an artifact of the step size.

## 5. Discussion

The presence of the long tails and diminished peaks in the DyMES (but not the SIR) curves demonstrates the importance of downward causation and of capturing microstate behavior in shaping the disease trajectories. All the DyMES model *I*(*t*) incidence curves display a long tail: disease incidence peaks early, then decreases at a slower rate compared to both its initial rise and the downward tail predicted by the SIR model. While it might seem counterintuitive that more downward causation would increase the long tail, flattening the curve is a well-documented epidemiological strategy [13]. Here, the preventative measures represented by *K* do not impact the ability of susceptibles to become infected, so the disease is still able to spread, albeit slowly, with the reduced transmission rate.

The curve-flattening phenomenon exhibited by DyMES is reflective of certain real-life epidemics, such as the COVID-19 pandemic [13]. Sheltering in place and social distancing are downward causation measures that affect the transmission rate rather than decreasing the number of susceptible individuals the same way a vaccine would. The decrease is steeper for higher values of *K*, consistent with less downward causation signaling weaker preventative measures.

The striking differences between the SIR and the *K* → ∞ limit predictions occur because the DyMES model takes averages across the distribution of the microvariable when calculating derivatives. For both models, the number of infected individuals must increase and then eventually decrease (as opposed to monotonically increasing), as we are not replenishing susceptibles or assuming that recovered individuals become susceptible to the disease. However, the two trajectories differ dramatically in all three cases, with the DyMES curves having much smaller peaks than the SIR curves.

Both Eqs. 2 and 5, when expanded, contain *n*^2^ terms multiplied by a negative constant, which are treated differently by the DyMES and SIR models. The SIR model is deterministic, so the model squares *n* directly wherever *n*^2^ appears. When calculating ⟨*f*_*t*_⟩, the DyMES model takes the expectation of *n*^2^, which is greater than ⟨*n*⟩^2^. As a result, at any time *t*, the DyMES derivative (*G*⟨*f*_*t*_⟩) will always be smaller than the analogous SIR derivative 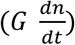 calculated using *m*_*t*_ − ⟨*n*⟩_*t*_ and ⟨*n*⟩_*t*_ in place of *s* and *n* in Eq. 2. At *t* = 0, the DyMES distribution is geometric with ⟨*n*⟩ = 1 and ⟨*n*^2^⟩ = 3, so the difference between *dn/dt* and ⟨*f*_*t*_⟩ is *c*_0_Var(*n*) = 2*c*_0_. For example, the difference is 0.7177 − 0.6931 = 0.0246 = 2 × 0.0123 in the high *c*_0_/high *d*_0_ case at time *t* = 0.

As *t* increases and the DyMES distribution diverges from the geometric with mean ⟨*n*⟩, the difference between ⟨*n*⟩^2^ and ⟨*n*^2^⟩ grows, and the DyMES derivative remains smaller than the corresponding SIR derivative before the SIR peak. Both models begin with *I*(0) = 100, but because the DyMES derivatives are less than the SIR derivatives when the SIR curve rises, the peak is lower for the DyMES models. In the high *c*_0_/high *d*_0_ and medium *c*_0_/medium *d*_0_ cases, the two curves intersect after the SIR peak because the SIR model depletes susceptibles faster than the DyMES model. In the low *c*_0_/low *d*_0_ case, the curves do not cross.

When we examine the log probability mass functions for the DyMES and the geometric distributions, we can distinguish the DyMES from the geometric distribution by the former’s positive second derivative. The curvature of the log geometric distribution is zero. Meanwhile, for the DyMES distribution, the curvature depends on *λ*_2_. The curvature is positive if *λ*_2_ is positive.

As mentioned above, expanding the transition function described in Eq. 5 reveals an *n*^2^ term that appears in the exponent of *P*(*n*|*I, dI/dt*) (Eq. 6), making the DyMES distribution’s functional form resemble that of a Gaussian. The *n*^2^ term is the highest degree term in Eq. 5, making the curvature independent of *n* and non-negative if *λ*_2_ is non-negative (Appendix B). *λ*_2_ = 0 at *t* = 0, so the DyMES distribution starts as a geometric distribution. Changing *I, K*, and *λ*_2_ introduces curvature into the log probability distributions that we see in Figure 2. Changing *c*_0_ and *d*_0_ also affects the curvature. If we are given the same values of *t, I, K*, and *λ*_2_, increasing *c*_0_ and *d*_0_ will increase the curvature, shifting more mass to the smaller values in the distribution, and vice versa.

The long tail and substantially lower peak value in overall incidence and the curvature in the log probability distributions are characteristic of the DyMES model. This model could be tested by gathering disease incidence data across many groups, each with many individuals. If we determine that the initial distribution of disease incidence resembles a geometric distribution, we can track the micro- and macrostates over time to examine any potential curvature in the log of the empirical probability distribution as well as any long tails appearing in the epidemic’s trajectory. This test would also allow determination of the value of *K*.

There exist two potential datasets: (1) serological survey data for low-pathogenicity avian influenza (LPAI) in 64 turkey farms [14] and (2) mortality and testing data for HPAI in poultry (chicken and turkey) across 72 commercial farms [15]. However, the first dataset only yields the final number of infected individuals, so we cannot test distribution evolution across time, and the second dataset breaks the assumption of homogeneity across groups, as turkeys have been shown to have higher transmissibility of HPAI [15].

The medium *c*_0_/medium *d*_0_ runs reveal a limitation of our DyMES model in epidemics with high *R*_0_. The basic reproduction number 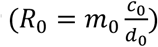 is highest for the medium *c*_0_/medium *d*_0_ simulations (*R*_0_ = 3.35), so there are the most expected new infections from a single infected individual. The model functions well through the initial increase and the peak. As total incidence falls, the infection still spreads in certain subgroups, as the proportion of the population unable to contract disease is less than the proportion necessary for herd immunity. For example, within our DyMES model, we could have many groups with low disease incidence but a few with higher disease incidence. This pattern of incidence shifts the probability distribution to have mass concentrated only at zero and the highest possible values (near *m*_*t*_), decreasing −*Gd*_0_⟨*n*⟩ and preventing satisfaction of the DyMES model’s constraints. Lower *K* decreases the effective *c*_0_ and therefore decreases *R*_0_ as *t* increases, so this phenomenon no longer occurs.

This paper describes only one way to model downward causation in an epidemic setting, and there are many other potential methods. Future studies could modify the transition function so that the downward causation term modifies the number of susceptible individuals or changes in response to the cumulative number of infected individuals. Just as the SEIATR model [5] builds on the SIR model, we could modify our DyMES model to introduce more complex mechanisms, such as the birth of new susceptibles, infection from asymptomatic individuals, vaccination that decreases the susceptible pool as well as the transmission rate, disease seasonality, and pathogen mutation. Future work could revisit the derivative constraint to model the full trajectory in epidemics with higher *R*_0_ and extend DyMES beyond a single microvariable–macrovariable pair to systems with two of each, extending the theory as well.

## Appendices

### Appendix A: Table of return times

**Table A.1:** Times taken to return to 100 for the DyMES finite *K* and DyMES *K* → ∞ limit models. Lower bounds exist where exact return times cannot be calculated due to constraints not being satisfied.

| Disease | DyMES $K=25$<br>time to return to<br>100 (days) | DyMES $K=100$<br>time to return to<br>100 (days) | DyMES $K=400$<br>time to return to<br>100 (days) | DyMES $K \rightarrow \infty$<br>limit time to<br>return to 100<br>(days) |
| --- | --- | --- | --- | --- |
| HPAI (high $c_0$/high $d_0$) | 93.98 | 65.14 | 26.54 | 21.46 |
| NDV (medium $c_0$/medium $d_0$) | 257.92 | 164.55 | > 61.53 | > 46.91 |
| FMD (low $c_0$/low $d_0$) | 246.85 | 185.18 | 83.46 | 63.85 |

### Appendix B: Derivation of curvature terms for DyMES and geometric distributions

Expanding Eq. 5 yields:

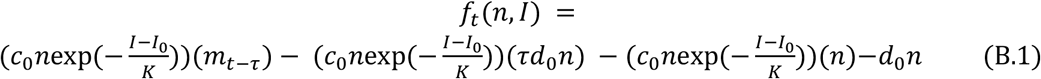

which then simplifies to:

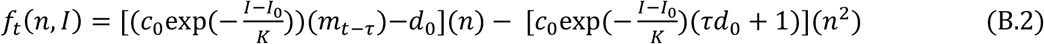

Substituting Eq. B.2 into Eq. 6 yields, for some normalization constant *z*:

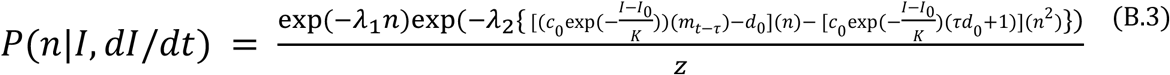

and

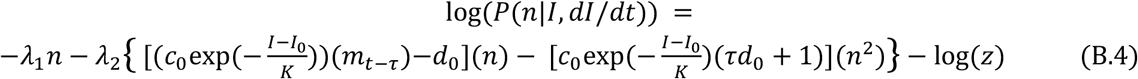

Taking the second derivative of Eq. B.4 with respect to *n* yields the curvature of the log DyMES distribution:

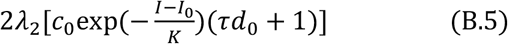

For the geometric distribution, we begin with:

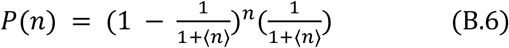

and

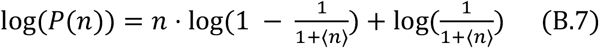

The second derivative of Eq. B.7 with respect to *n* is zero, which means there is no curvature in the log probability distribution.

### Appendix C: Derivation of failure condition for DyMES distribution

We begin with Eq. 5. The infection term *f*_*t*_(*n, I*)+ *d n* is non-negative from 0 to 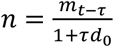.

Therefore:

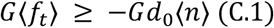

with equality at *n* = 0 and 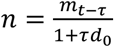.

Following from Eqs. 3 and 4, as well as the above:

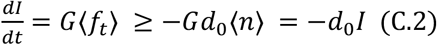

When *I* = *G*⟨*n*⟩ but Eq. C.2 above does not hold, then no distribution satisfies both constraints and the DyMES model has no solution.

## Author Contributions

**Meilin Yen:** Conceptualization, Methodology, Software, Validation, Formal analysis, Writing – original draft, Writing – review and editing, Visualization; **Pranav Muralikrishnan:** Conceptualization, Methodology, Software, Writing – review and editing; **John Harte:** Conceptualization, Methodology, Validation, Writing – original draft, Writing – review and editing, Supervision

## Code/Data Availability

All code can be found in the following GitHub repo: https://github.com/mmyen/DyMES-disease-transmission.

## Declaration of generative AI and AI-assisted technologies in the manuscript preparation process

During the preparation of this work the authors used Claude Code to help generate code for producing figures. After using Claude Code, the authors reviewed and edited the content as needed and take full responsibility for the content of the published article.

## Acknowledgements

We thank Keon Abedi, Amber He Wei, Kayli Huang, and Alex Landry for useful discussions and comments on earlier drafts.

